# EvoMining reveals the origin and fate of natural products biosynthetic enzymes

**DOI:** 10.1101/482273

**Authors:** Nelly Sélem-Mojica, César Aguilar, Karina Gutiérrez-García, Christian E. Martínez-Guerrero, Francisco Barona-Gómez

## Abstract

Natural products, or specialized metabolites, are important for medicine and agriculture alike, as well as for the fitness of the organisms that produce them. Microbial genome mining aims at extracting metabolic information from genomes of microbes presumed to produce these compounds. Typically, canonical enzyme sequences from known biosynthetic systems are identified after sequence similarity searches. Despite this being an efficient process the likelihood of identifying truly novel biosynthetic systems is low. To overcome this limitation we previously introduced EvoMining, a genome mining approach that incorporates evolutionary principles. Here, we release and use our latest version of EvoMining, which includes novel visualization features and customizable databases, to analyze 42 central metabolic enzyme families conserved throughout Actinobacteria, Cyanobacteria, Pseudomonas and Archaea. We found that expansion-and-recruitment profiles of these enzyme families are lineage specific, opening a new metabolic space related to ‘shell’ enzymes, which have been overlooked to date. As a case study of canonical shell enzymes, we characterized the expansion and recruitment of glutamate dehydrogenase and acetolactate synthase into scytonemin biosynthesis, and into other central metabolic pathways driving microbial adaptive evolution. By defining the origins and fates of metabolic enzymes, EvoMining not only complements traditional genome mining approaches as an unbiased and rule-independent strategy, but it opens the door to gain insights into the evolution of natural products biosynthesis. We anticipate that EvoMining will be broadly used for metabolic evolutionary studies, and to generate genome-mining predictions leading to unprecedented chemical scaffolds and new antibiotics.

**DATA SUMMARY:** Databases have been deposited at Zenodo; DOI: 10.5281/zenodo.1162336 http://zenodo.org/deposit/1219709

Trees and metadata have been deposited in MicroReact

GDH Actinobacteria https://microreact.org/project/r1IhjVm6X

GDH Cyanobacteria https://microreact.org/project/HyjYUN7pQ)

GDH Pseudomonas https://microreact.org/project/rJPC4EQa7

GDH Archaea https://microreact.org/project/ByUcvNmaX

ALS Cyanobacteria https://microreact.org/project/B11HkUtdm

EvoMining code has been deposited in gitHub https://github/nselem/evomining

Docker container in Dockerhub https://hub.docker.com/r/nselem/evomining/

**We confirm all supporting data, code and protocols have been provided within the article or through supplementary data files.**

**IMPACT STATEMENT:** EvoMining allows studying expansion-and-recruitment events of enzyme families in prokaryotic lineages, with the goal of providing both evolutionary insights and a genome mining approach for the discovery of truly novel natural products biosynthetic gene clusters. Thus, by better understanding the origin and fate of gene copies within enzyme families, this work contributes towards the identification of lineage-dependent enzymes that we call ‘shell’ enzymes, which are ideal beacons to unveil ‘chemical dark matter’. We show that enzyme functionality is a continuum, including transition enzymes located between central and specialized metabolism. To exemplify these evolutionary dynamics, we focused in the genes directing the synthesis of the sunscreen peptide scytonemin, as the two key enzymes of this biosynthetic pathway behave as shell enzymes and were correctly identified by EvoMining. We also show how evolutionary approaches are better suited to study unexplored lineages, such as those belonging to the Archaea domain, which is systematically mined here for novel natural products for the first time. The release of EvoMining as a stand-alone tool will allow researchers to explore its own enzyme families of interest, within their own genomic lineages of expertise, by taking into account the lessons learned from this work

## INTRODUCTION

Natural products (NP), or specialized metabolites, are naturally occurring molecules widely used in medicine and in other applications (1). NP are typically encoded in biosynthetic gene clusters (BGC) found in the genomes of a wide range of organisms. From sources as diverse as bacteria, fungi and plants, there are around 1800 NP with their cognate BGC experimentally characterized. This body of knowledge, contained in a community-driven hierarchical repository called the Minimum Information about a Biosynthetic Gene cluster (MIBiG) (2), allows investigating newly sequenced microbial genomes as never before. Indeed, current availability of around half a million prokaryotic genomes in public databases has not only invigorated research into NP, but it has also prompted the development of novel genome mining bioinformatic tools (3,4). The latter have evolved from simple sequence similarity searches of known biosynthetic enzymes, with an emphasis in the domains of Polyketide Synthases (PKS) and Non-Ribosomal Peptide Synthetases (NRPS)(5,6); to genome-mining platforms that look into complete BGC, such as the antibiotics and Secondary Metabolite Analysis shell (antiSMASH) (7). For instance, only in closed genomes, a tantalizing total number of 32,584 BGCs have been identified using antiSMASH (8).

Even if it is safe to state that known classes of NP BGC are successfully predicted by antiSMASH, it is worthwhile to note that not all BGC correspond with well-known biosynthetic enzyme classes. Only within MIBiG there are 231 BGC (12.7%) classified as “Other”, which indeed lack a PKS, NRPS or any of the other enzyme classes characteristic of specialized metabolism. Absence of known biosynthetic enzymes turns these BGC into ‘atypical’, hard to identify, relating them to the term ‘chemical dark matter’ (9). An example of this scenario is provided by the BGC of the cyanobacterial sunscreen scytonemin (10), which includes ScyB and ScyA as the two key biosynthetic enzymes sustaining the synthesis of this specialized metabolite (11,12). Interestingly, ScyB and ScyA are distant homologues of glutamate dehydrogenase (GDH) and acetolactate synthase (ALS), respectively, which take part in the reversible oxidative deamination of glutamate to a-ketoglurate and ammonia (13) and in the synthesis of branched-chain amino acids (14).

It is clear therefore that despite the overwhelming amount of accurately predicted BGC, there is still plenty of space to discover and prioritize novel biosynthetic systems. Only in the genus *Streptomyces* this has been estimated between 15% and 26% BGC per genome, which are actually different from those identified by antiSMASH (15). To address the problem of the limited novelty revealed by genome mining approaches based in sequence similarity searches, we have previously introduced the use of evolutionary principles driving the emergence of NP BGC (15,16). The latter gave place to EvoMining, which recapitulates enzyme evolutionary events as follows: any given enzyme family (EF) may undergo expansions (17) due to gene duplication, leading to paralogues, and/or horizontal gene transfer (HGT), leading to xenologues. The emerging extra gene copies may be retained in the genome when they provide an advantage, as they evolve into novel enzyme functions (18) to serve as raw material for new metabolic pathways. Since genes involved in a metabolic pathway tend to cluster together in bacterial genomes, conserved genomic vicinity can be taken as an indication of related gene functionality (19), whereas phylogenetic reconstruction of these EF can differentiate between conserved copies devoted to central metabolism, from expansions recruited into NP biosynthesis or other metabolic adaptations.

To date, EvoMining has been used to show the occurrence of conserved EF that fulfill related biochemical functions but yet with different physiological roles (16,20). It has also been used for the discovery of NP BGC that include unprecedented biosynthetic enzymes essential for the synthesis of arsenolipids in *Streptomyces* (15). As a related but independent follow-up of EvoMining, we have very recently released a phlyogenomic approach, termed CORe Analysis of Syntenic Orthologs to prioritize Natural products BGC, or CORASON. This algorithm addresses the evolutionary relationships between BGC, allowing to comprehensively identifying all genomic vicinities in which particular biosynthetic gene cassettes are found (21). Based in similar evolutionary ideas to those embraced by EvoMining, the Antibiotic Resistance Target Seeker, or ARTS (22) exploits the fact that some antibiotics function by interfering central metabolic enzymes, and therefore antibiotic-producing bacteria have mechanisms of self-protection encoded in extra gene copies. Although examples of the discovery of novel biosynthetic systems using CORASON or ARTS remain to be reported, these approaches together with EvoMining add to the growing notion that evolutionary paradigms can aid in the discovery of antibiotics (23).

Although elegant in their simplicity, the abovementioned evolutionary ideas oversimplify a far more complex scenario in which different evolutionary histories can involve different metabolic origins and fates (24). For instance, the boundaries of central metabolism are hard to define, as conserved or core enzymes of genomic lineages tend to differ broadly, even within closely related clades or organisms (25). Additionally, not all extra copies of expanded enzymes are recruited into specialized metabolism, as previously highlighted as a criticism of EvoMining (3). Some of the expanded EF may remain in central metabolism providing certain level of metabolic redundancy, as it is the case of pyruvate kinases in glycolysis (20) or ketol-acid reductoisomersases in the biosynthesis of branched-chain amino acids (26) in *Streptomyces* species. Alternatively, expanded EF could serve other physiological roles related to morphological development (27) or may represent metabolic adaptations that involve the use of different cofactors as in glutamate dehydrogenase (GDH) of Archaea species (28). Although EvoMining overcomes the latter caveats by prioritizing extra copies that are similar in sequence to enzymes from NP BGC with experimental support, there is much to be understood about the evolution of enzymes during the assembly of BGC directing the synthesis of NP.

An open question along these lines is whether we can identify novel enzymes that define a completely new class of BGC, rather than only identifying accessory or precursor-supply enzymes. To address this question, here we developed EvoMining to allow customization of its databases. We then performed a systematic analysis of expansion-and-recruitment events in different prokaryotic lineages, including divergent taxa (Actinobacteria, Cyanobacteria, *Pseudomonas* and Archaea). Selected results were visualized with CORASON (21), as a feature that can be integrated into EvoMining, depicting the evolutionary dynamics leading to new BGC. We analyzed, as a case study, the BGC of scytonemin, a pigment exclusively produced by Cyanobacteria (10). Our results directed us to rethink EvoMining to incorporate ‘shell’ enzymes, which are defined as orthologues shared by the majority, but not all genomes, of any given taxonomic group (29). We demonstrate that this class of enzymes has great potential for the discovery of novel NP BGC, which agrees with the result that EF recruitments are taxa-dependent, better understanding the origin and fate of metabolic enzymes.

## METHODS

### EvoMining container

EvoMining version 2.0 was developed as a standalone comparative genome-mining tool using Perl as coding language and docker (30) as packaging platform. Previous recommendations for the use of containers were adopted (31). A simplified version of CORASON code, reported simultaneously (21), was included within EvoMining 2.0 container to allow visualization of the genomic vicinity. EvoMining dependencies including BLAST, MUSCLE, Gblocks, FastTree, as well as Newick utilities, were wrapped in the EvoMining docker container, available at the DockerHub nselem/evomining. EvoMining 2.0 operational details can be consulted in the user manual available at https://github.com/nselem/evomining/wiki. The increased performances obtained by these developments, including the biological insights that can be gained, are summarized in **Table 1**.

**Table 1.** EvoMining 2.0 novel developments and associated biological insights.

|  | EvoMining 1.0 | EvoMining 2.0 | Main results |
| --- | --- | --- | --- |
| **Tool** | Consulting website | Stand-alone tool in docker container. | Users can visualize EF using a colour code according to their metabolic origin and fate. |
| **Databases** | Fixed | Provided by the user | Databases are customizable. |
| **Genome-DB** | Actinobacteria (230 genomes) | Actinobacteria (1244 genomes)<br>Cyanobacteria (416 genomes)<br>*Pseudomonas* (219 genomes)<br>Archaea (876 genomes) | Expansion rate is related to genome size. The total number of expansions is similar across lineages until a genome size of 5 Mbp. After this threshold *Pseudomonas* surpasses Actinobacteria, which in turn exceeds Cyanobacteria. There are no reported archaeal genomes in this size range, **Fig 2b**. |
| **Enzyme-DB** | Actinobacteria (106 families) | Comparative framework consisting of 42 conserved EF | Construction on the Enzyme DB by identification of 42 EF conserved throughout Actinobacteria, Cyanobacteria, *Pseudomonas* and Archaea, **Fig 2a**. Expansions, recruitments and phylogenetic histories is lineage-dependent, **Fig 3a**.<br>Families in Enzyme DB are often part of the shell genome, i.e. they are shared by the majority, but not all, of the genomes in a lineage, **Fig 3b,c**. Shell EF show expansion- and recruitment events. These two observations stand behind current EvoMining paradigm, leading to the use of core metabolism and shell enzymes as key concepts. |
| **NP-DB** | Manual curation (226 BGC) | MIBiG (1296 BGC) | BGC from different taxa, allowing to track the fate of metabolic enzymes within vertical lineages, but also after horizontal gene transfer. **Fig 3**. |

### EvoMining expansion-and-recruitment algorithm

Expansion-and-recruitment events of EF represent the first output of EvoMining. Expanded EF were obtained using BLASTp between seed enzymes of an EF (Enzyme DB) against a database of genomes (Genome DB), using an e-value of 0.001 and a score of 100. Conserved sequences within the expanded EF were identified after bidirectional best hits (BBH) against the Enzyme DB. Expansions were defined as previously (15,16), and this information was presented as a heat plot, **Fig. S1.** This plot pinpoints, for each EF, the organisms that have extra enzyme copies. Events of enzyme recruitment into specialized metabolism were determined by BLASTp searches with an e-value of 0.001, where every sequence in the expanded EF were queried against a DB of genes that encode for NP biosynthetic enzymes (NP DB) (2). These actions were systematized as part of the EvoMining algorithm.

### EvoMining phylogenetic reconstruction-and-visualization algorithm

EF found to exhibit expansion-and-recruitment events were aligned with MUSCLE v3.2 (32) and automatically curated with Gblocks v0.91b (33). Parameters used included five positions as minimum block length and ten as the maximum number of contiguous nonconserved positions. Positions with a gap in more than 50% of the sequences were filtered and were not used for the final alignment. Curated alignments were phylogenetically reconstructed with FastTree 2.1 (34), which is an approximately maximum likelihood method. These actions, leading to EvoMining trees per EF, were systematized as part of the EvoMining algorithm.

EvoMining trees provide evolutionary insights into the metabolic origin and fate of members of any given EF, by differentiating gene copies through a color-labeling process, **Fig. 1**. The most conserved sequences are catalogued as coming from central metabolism, and these sequences are marked in red. Cyan color is used for depicting sequences that are part of a BGC, as revealed by antiSMASH (7). Although EvoMining does not calculate by default antiSMASH predictions these can be provided as indicated in the user’s manual. Purple is used for depicting transition enzymes, defined as those sequences that are simultaneously identified by antiSMASH, but which seem to come from central metabolism, whereas gray is used to highlight extra copies with an unknown metabolic fate, which may include other metabolic adaptations as previously shown in S. *coelicolor* (20). Finally, blue color is used to indicate the few recruitment events included within MIBiG (2), whereas green is used to denote EvoMining predictions, which represent the outputs with the largest potential to unveil unprecedented biosynthetic enzymes and their pathways. Tree labeling with a color code was automatized by the Newick utilities (35).

**Figure 1.**
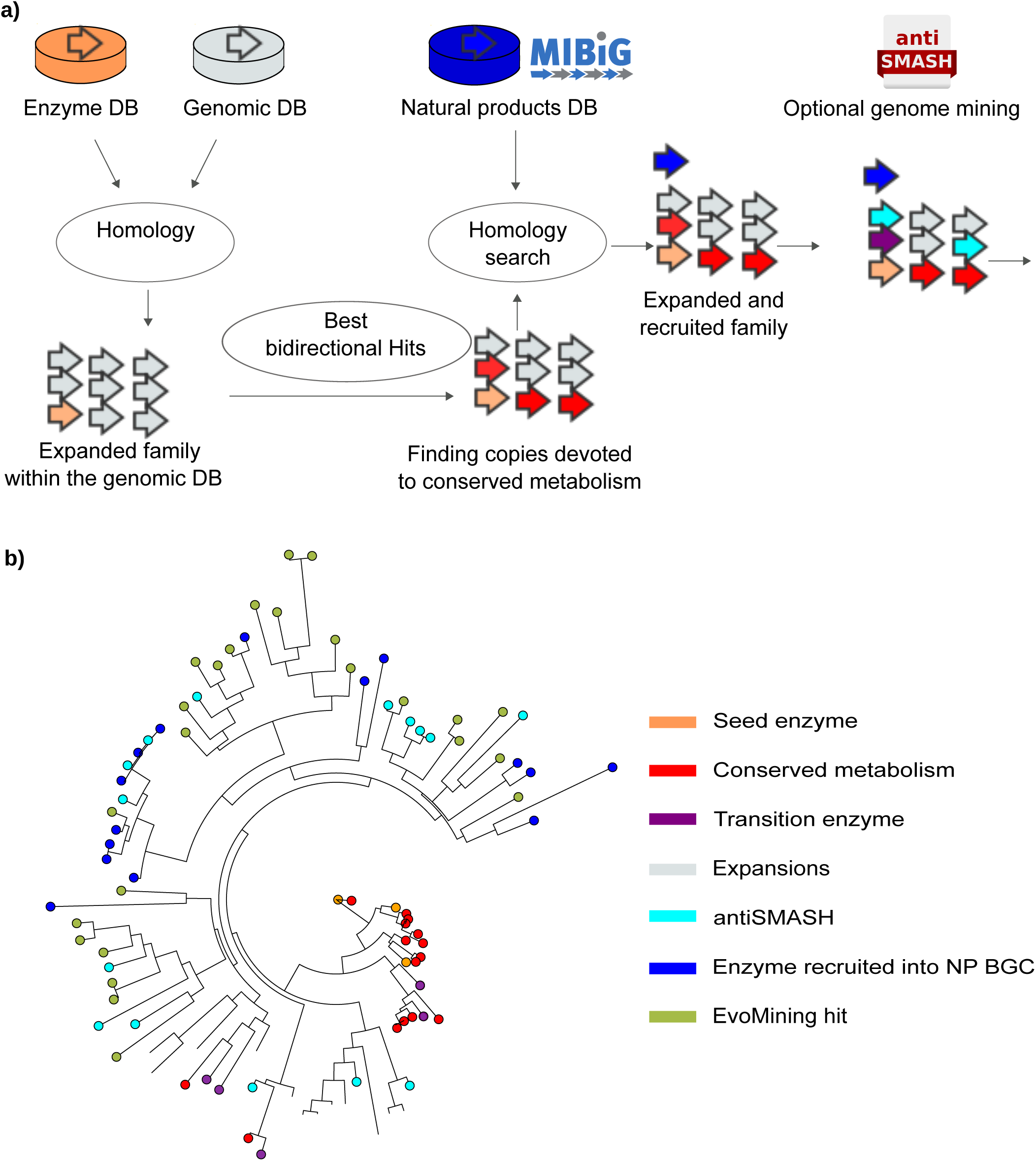
EvoMining pipeline, databases and enzyme’s origin and fate. **a)** EvoMining pipeline and databases. Homologues and expansions of seed enzymes (orange) from Enzyme DB are searched in the Genome DB. The outcome is integrated as the EF. BBHs of seed enzymes (red) are marked as conserved metabolism. The EF are amplified after being compared against the NP DB (blue) to find enzymes of the family recruited into NP biosynthesis. Optionally, antiSMASH predictions (cyan) can be added by the user. Enzyme predictions belonging to specialized metabolism by antiSMASH, but that at the same time were marked red as members of conserved metabolism, are defined as transition enzymes (purple). **b) EvoMining phylogenetic output**. A phylogenetic reconstruction of an EF is done to differentiate expansions of unknown metabolic fate (gray) from enzymes closer to recruitments (blue) or to conserved metabolic enzymes (red). The resulting enzymes are the EvoMining hits (green), which represent novel enzymes devoted to specialized metabolism.

Enriched metadata, such as gene copy number by organism and functional information provided by the platform Rapid Annotation using Subsystem Technology, or RAST (36), was also added. Trees and metadata were arranged such that they are compatible with the specialized visualization tool Microreact (37).

### EvoMining Databases

Three databases are needed to run EvoMining, the Genome DB, the Enzyme DB and the NP DB (or MIBiG). These databases are provided as starting parameters through the command line before EvoMining is run. The integration of the current versions of these databases is described as follows, **Table 1, Fig S2.**

*Genome DB*. Previous EvoMining Genome DB comprised 230 Actinobacteria genomes, including 50 different genera. In EvoMining 2.0 the Actinobacteria Genome DB was expanded to 1245 genomes, including 193 genera. Additionally, three new Genome DB were constructed and integrated, including organisms belonging to Cyanobacteria (416 genomes), *Pseudomonas* (219 genomes) and Archaea (876 genomes).

Since EvoMining predictions are based in the ability of its algorithm to identify expanded enzymes, and not complete BGC, draft genomes with an average of five genes by contig were included. The selected genome sequences were retrieved from public databases (NCBI), as available in January of 2017, and functionally annotated by RAST (36) and antiSMASH (7) with a parameter cf_threshold of 0.7. To identify antiSMASH predictions in the final tree these results were delivered into an internal database, called antiSMASH DB.

*Enzyme DB*. Previous EvoMining Enzyme DB comprised 106 EF, which were selected in the basis of metabolic criteria, namely, enzymes from central metabolic pathways that could be unambiguously annotated within the genome-scale metabolic models of *Streptomyces coelicolor, Mycobacterium tuberculosis* and *Corynebacterium glutamicum* (15). These 106 EF comprised 339 actinobacterial amino acid sequences, used as seeds for our previous proof-of-concept analyses. In this version, the newly created Enzyme DB consists of a common set of central EF identified in at least one seed genome from Cyanobacteria, *Pseudomonas* and Archaea. To avoid missing hits due to gaps in sequences the seed genomes providing query enzymes were selected as they are contained in one single contig. For Cyanobacteria, these genomes were those from *Cyanothece sp*. ATCC 51142, *Synechococcus sp*. PCC 7002 and *Synechocystis sp*. PCC 6803; for the genus *Pseudomonas*, those from *P. fluorescens* pf0-1, *P. protegens* Pf5, *P. syringae* and *P. fulva* 12-X; and for the domain Archaea, those from *Natronomonas pharaonis, Methanosarcina acetivorans, Sulfolobus solfataricus* and *Nanoarchaeum equitans* Kin4-M. Seed sequences were determined after BBH from the previous Actinobacteria Enzyme DB (15), using Metaphor tool (38). Default parameters were used to discard hits that do not account for at least 30% of sequence identity across an alignment length of 80% of the two protein sequences. The original 106 actinobacterial EF was filtered to 42 EF, which are shared by the seed genomes of Actinobacteria, Cyanobacteria, *Pseudomonas* and Archaea.

*NP-DB*. The original EvoMining NP DB included 226 manually curated BGC (15). In this work the NP DB is that of MIBiG (2), which in its release of August 2018 as MIBiG v1.4, includes 1,813 NP BGC and a total of 31,023 protein sequences.

### EvoMining analysis of EF from scytonemin BGC

After EvoMining analysis of the conserved 42 EF, we focused in those present in the scytonemin BGC. Detailed analyses of this BGC included phylogenomic reconstruction and genomic vicinity visualization using CORASON (21). Details of these EF, including their distribution, **Table S1 & Fig. S4**, and expansion trees, **Fig. S5-S11**, are provided as supplementary information. Based in the results obtained by EvoMining, chemical diversity of the scytonemin BGC was predicted utilizing their conserved enzyme repertoire, as revealed by CORASON, and the domain organization of the NRPS and NRPS-PKS assembly lines, using antiSMASH 3.0 (7) and PKS and NRPS analysis (6). Up to 30 genes upstream and downstream the *scyA* gene were retrieved and analyzed. For CORASON analysis the amino acid sequences of ScyA and ScyB were concatenated and aligned using Muscle v3.2 (32). A phylogenetic reconstruction was produced from the amino acid alignment matrix using MrBayes v3.2 (39) with a gamma distribution type range and 1 million generations. ScyA and ScyB sequences from *Scytonema tolypothrichoides* VB 61278 JXCA01 were used as outgroup.

## RESULTS & DISCUSSION

### Increased applicability of EvoMining

To transform the website EvoMining version 1.0 into a genome-mining tool that allows analysis and visualization of large genomic datasets, we first aimed at customizing its databases, **Table 1**. As a result, the three EvoMining inputs, (i) Genome DB, (ii) Enzyme DB (originally called PSCP, from Precursor Supply Central Pathways) and (iii) NPs DB, can be modified, replaced or expanded by the user, **Fig 1a**. For each EF, the pipeline produces an interactive, color-coded tree of the expanded EF, **Fig 1b**. Colors in the tree show information about the metabolic origin and/or fate of homologous enzymes present in the Genome DB. As described in methods, red stands for central metabolism, which can be orange if it coincides with the seed enzyme; purple is used for transition enzymes; gray for expansions of unknown origin and/or fate; cyan for antiSMASH predictions; blue for MIBiG recruitments; and green for EvoMining predictions. Visualization of the genomic vicinities in which each of these enzymes is encoded can be generated. Thus, the newly incorporated changes allow, at a glance, to explore the evolutionary dynamics of EF, from central to different forms of specialized metabolism, with an emphasis on NP biosynthesis. Novel capabilities and potential insights provided by EvoMining are summarized in **Table 1**, and the derived insights are developed as follows.

To exploit these improvements, we first analyzed whether expansion-and-recruitment events of EF are lineage-dependent, and how this may relate to genome size. The concept of shell enzymes and its potential to unveil novel pathways is then presented. As further discussed in the following sections, we defined core EF are those with a copy in every genome of a lineage, whereas shell EF are defined as those enzymes that are shared by more than 50% of the genomes of any given genomic lineage. The presented analyzes were based in 42 EF conserved in the seed genomes of the four lineages considered in this study. Second, scytonemin BGC is characterized as an example of the potential and intrinsic features of EvoMining algorithm. The selection of this BGC obeys to the fact that it is composed by atypical biosynthetic enzymes that happened to be included within the 42 EF analyzed, namely, GDH and ALS. These EF were detected by EvoMining as originated in central metabolism, with their fate in specialized metabolism or central metabolic pathways related to adaptive microbial physiologies involving different cofactors, depending on the taxon analyzed.

### Enzyme expansion rates are lineage-dependent

To further gain insights into the evolution of enzymes and the pathways in which they take part, we exploited the taxonomic coverage of the selected lineages. The newly assembled Genome DB consisted of the phyla Actinobacteria and Cyanobacteria, the genus *Pseudomonas*, and the domain Archaea. Results shown in related figures are always presented following this order. The selection of these taxa obeyed to the possibility of analyzing both well-known NP-producing microorganisms, namely, Actinobacteria (602 MIBiG BGC), Cyanobacteria (60 MIBiG BGC) and *Pseudomonas* (53 MIBiG BGC); but also poor NP-producing taxa, such as Archaea (0 BGC in MIBiG version 1.3), which represents a domain whose NP biosynthetic capabilities have not been investigated until very recently (40). Based in these Genome DB, and following the scheme of **Fig. 2a**, a set of EF shared among them was first identified. Notably, only a fraction of the original 106 actinobacterial EF was found conserved as new taxa was incorporated. Thus, each taxon-specific DB contains only 42 EF, **Table S2**. The observation that 64 EF are not conserved throughout these four taxa reflects on the species or lineage specificity of microbial metabolism (41), an intrinsic feature that is acknowledged and better exploited in this new version of EvoMining.

**Figure 2.**
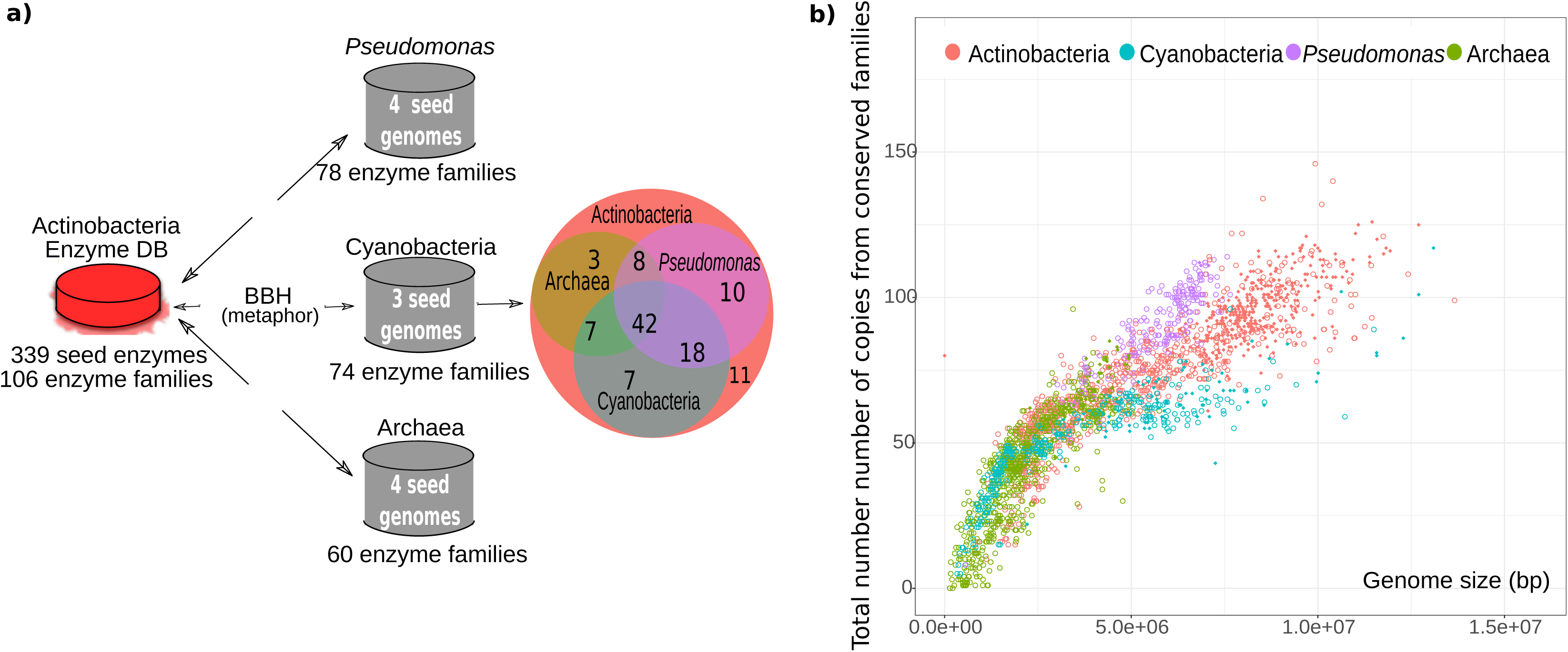
EvoMining Enzyme DB. **a)** Previous EvoMining Enzyme DB was filtered to establish a common set of 42 conserved enzyme families for the phyla Actinobacteria and Cyanobacteria, the genus *Pseudomonas* and the domain Archaea. b) All taxa show expansions of conserved EF and these expansions correlate with genome size. Differences in expansion rates across taxa are principally noted after a genome size greater than 5 Mbp. At this threshold *Pseudomonas* surpasses Actinobacteria expansions which in turn overcomes Cyanobacteria. No bigger Archaeal genomes were reported.

Using these conserved 42 EF we found that in all lineages the expansion rates behave similar until a genome size of 5 Mbp. After this threshold the total number of sequences in the 42 EF grows faster in the genus *Pseudomonas* than in the phylum Actinobacteria, which in turn surpasses the phylum Cyanobacteria and the domain Archaea, **Fig. 2b**. The latter observation may be related to the fact that there are no reports of Archaea genomes with sizes comparable to those typically found in *Streptomyces* or *Pseudomonas* (>5 Mbp). In contrast, Cyanobacteria, despite having big genomes, were the taxon with the fewer expansions. This could be a real biological observation, or that the selected Enzyme DB, by chance, lacks expansions in this lineage. In any case, these results suggest that EvoMining is better suited to analyze large genomes when used as a tool to generate NP BGC predictions.

Our results also show that the taxonomic orders with the greatest number of copies in the common Enzyme DB were *Streptomycetales* and *Nostocales*, in Actinobacteria and Cyanobacteria, respectively. This observation coincides with the fact that these two orders have the biggest genome size within their corresponding lineages, and these orders are well known to have metabolic diversity and biosynthetic potential. Interestingly, the class Halobacteria showed the largest number of expansions in Archaea, even though it is not the class with the biggest genome size in average, **Fig S3.**

Yet, this result is in agreement with the observation that archaeocines, diketopiperazines, carotenoids and other NP from Archaea have all of them being isolated from Halobacteria species, even though their NP BGC remained to be discovered (40). Thus, EvoMining is well suited to genome mine unexplored lineages with the potential to encode truly novel pathways.

Overall, these results show that expansions from central families correlate with genome size. However, the increment of the expansion rates is different in each genomic group, and this increment is not linear, **Fig 2b**. These results are important to direct the use of EvoMining, first, as they emphasize the importance of properly assembled and ad hoc Genome DB; and second, to better understand the predictions derived from its use when more than one genomic lineage, with different genome sizes, is explored. These points are revisited in the following subsections.

### EvoMining reveals the occurrence of extra copies in ‘shell’ enzymes

Having shown that divergent genomes experience different lineage-specific expansion rates, we then focused in the expansion-and-recruitment patterns across the different taxa, **Fig 3a**. Our analyses show that *Pseudomonas* has on average more copies per genome than any other taxa, as 54.8% of the 42 EF showed a maximum average copy number for this lineage. In contrast, Actinobacteria showed a maximum average in only 26.2% of the EF, while Archaea and Cyanobacteria had a similar result, as little as 9.5%, **Table S2**. While there are families like acetylornithine aminotransferase or ALS that are highly expanded in every lineage (coordinates A1 and E1, **Fig. 3a**), many others exhibit differential behavior. Such is the case of fumarate reductase iron-sulfur subunit (coordinate C3, **Fig. 3b**), which is highly expanded in Actinobacteria but has in average less than one copy per genome in Cyanobacteria.

**Figure 3.**
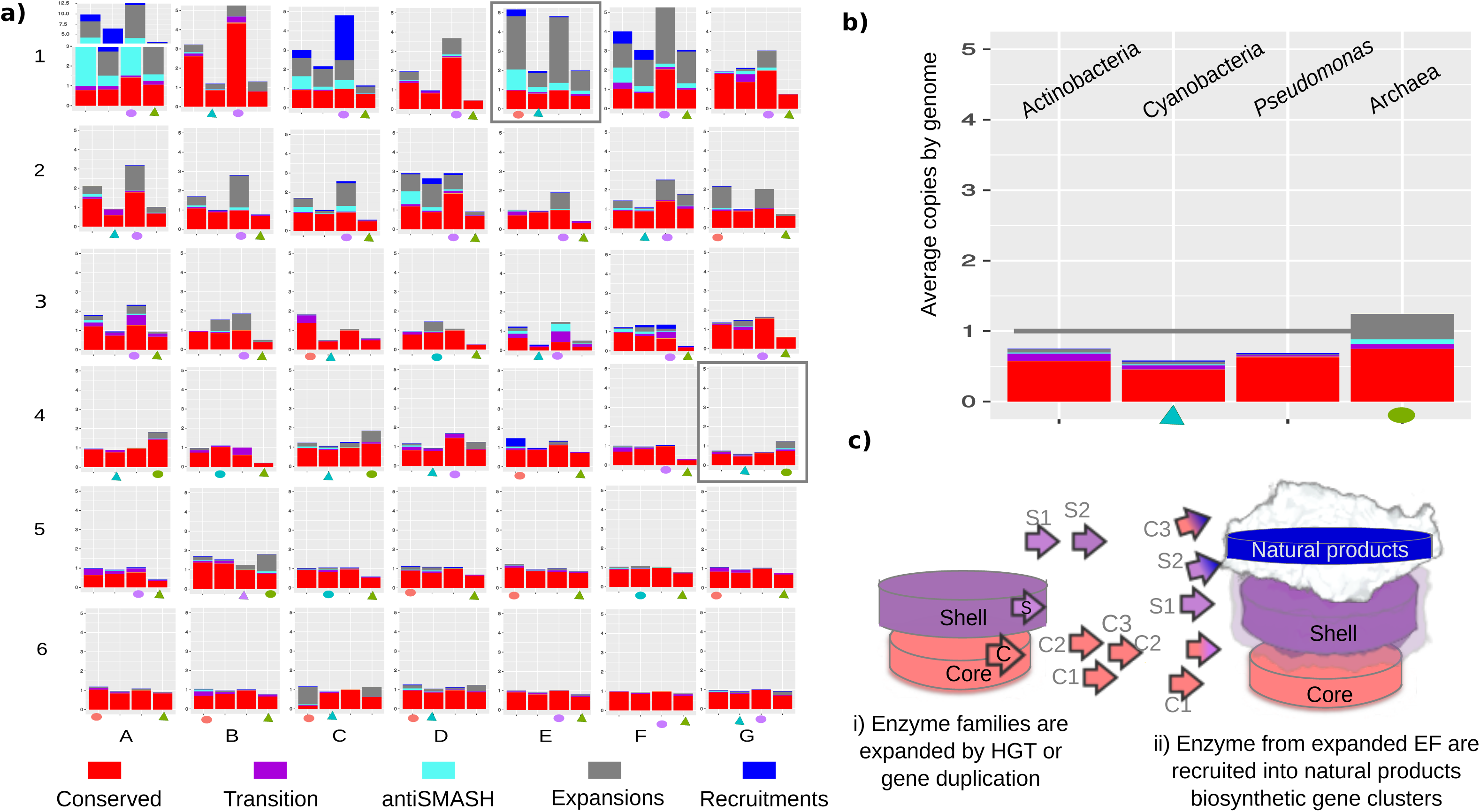
EvoMining profiles of conserved enzymes in selected genomic lineages. **a)** Expansions of the 42 conserved EF are colored as follows: red for conserved metabolism, blue for recruitments annotated at MIBiG, cyan for antiSMASH predictions of specialized metabolism, purple for the intersection between conserved metabolism and antiSMASH predictions, and gray for expansions without known metabolic fate. Triangles indicate the lineage with the largest number of copies by genome on average, and circles stands for the least expanded. Although Archaea tends to be the least expanded taxa this tendency reverts in families A4, C4, G4 (GDH) and B5. GDH and ALS in E1, in a box, are the origin of recruitments into scytonemin BGC. **b)** Zoom-in of G4 family showing that on average there is less than one copy by genome in Actinobacteria, *Pseudomonas* and Cyanobacteria, and consequently GDH is not in the core but in the shell genome of these lineages. **c)** Model for the appearance of shell enzymes, which are being expanded and recruited into NP BGC in the observed lineages. The definition of central metabolism for EvoMining purposes can be improved as conserved metabolism comprising both shell and core enzymes.

After inspecting these results in more detail it was interesting to note that some EF are not expanded, suggesting that even when the 42 EF are conserved throughout the four seed genomes selected for each taxa investigated, there are organisms that lack certain enzymes and most likely their metabolic pathways. Yet, the independent expansion levels of each EF, in general, showed the same pattern as that recorded overall, with *Pseudomonas* as the lineage with the largest number of expansions, and Archaea with the least expansions. However, there are certain families that do not follow this trend. For instance, GDH is one of the four EF with the largest expansion rate in Archaea. In fact, GDH has less than one copy per genome in the other three taxa, to the point that this EF is not part of the core of these lineages. Based in these observations therefore we define GDH as a member of the shell genome (29) of Actinobacteria, Cyanobacteria and *Pseudomonas*, consistent with the fact that it is not conserved but yet present in more than 50% of the genomes of each lineage, **Fig 3b**.

Specifically, for the GDH EvoMining results we found that antiSMASH predictions are present in Actinobacteria, Cyanobacteria and Archaea, but not in *Pseudomonas*. This observation coincides with the fact that the recruitment of GDH into a NP BGC, namely, that of scytonemin but also of the unrelated polyketide pactamycin (42), was only recorded for the former three taxa and not for *Pseudomonas* (see **Fig 4** in following subsection). These results together suggest that the evolution of specialized metabolism is lineage-dependent, but more importantly, that shell enzymes just as core EF, possess the potential to drive the evolution of NP BGC, **Fig. 3c**. The latter is a fundamental consideration that we previously overlooked when exploiting EvoMining as a genome mining tool to generate novel biosynthetic predictions (21).

**Figure 4.**
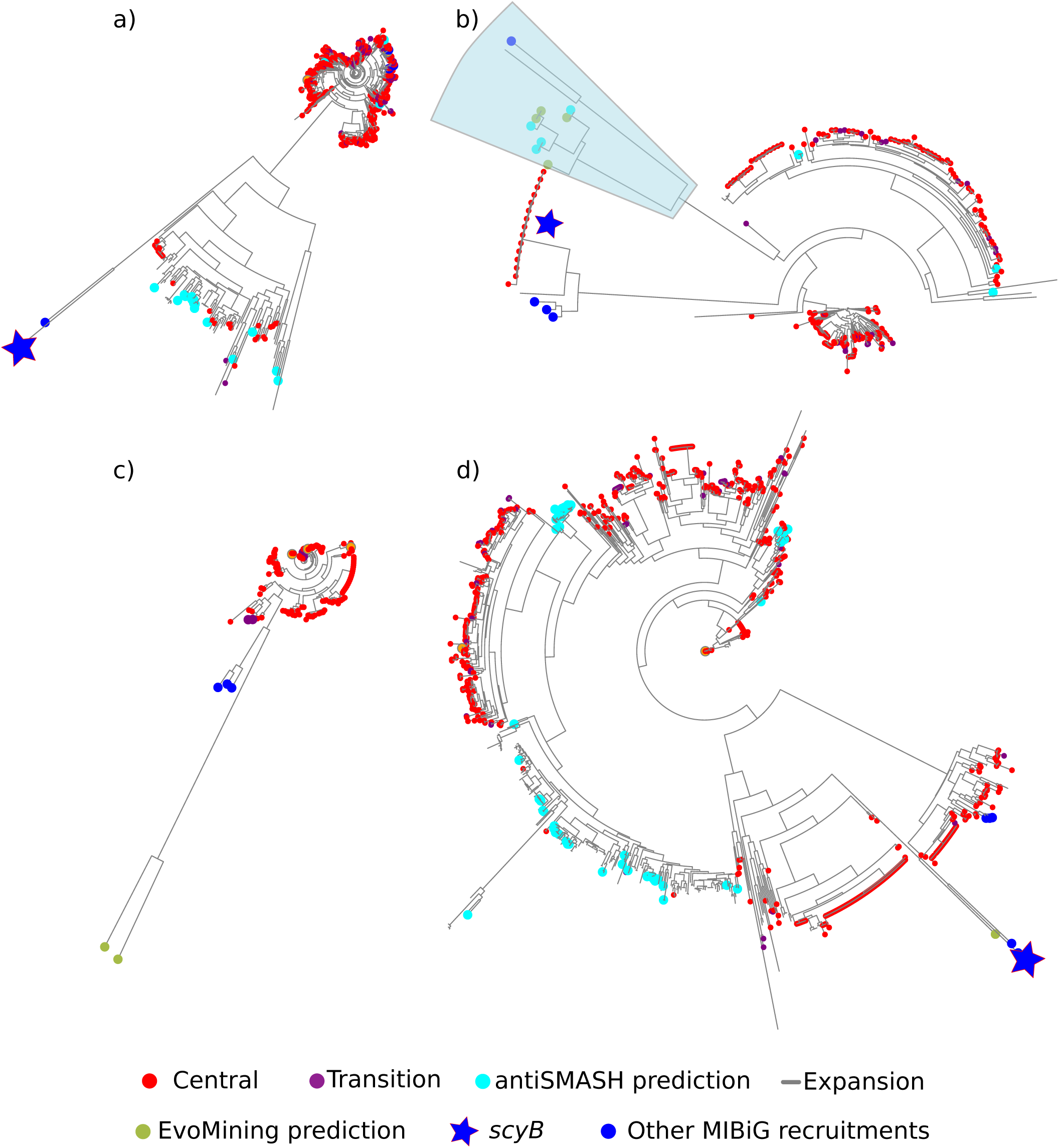
GDH EvoMining trees by taxa. **a)** Phylogenetic reconstructions reflect differences in expansion patterns in Actinobacteria, Cyanobacteria, *Pseudomonas* and Archaea. Actinobacteria has no EvoMining predictions, as its main expansion branch lacks MIBIG recruitments. Nevertheless, it is still possible that specialized metabolism may occur within those copies of unknown fate (gray). **b)** Cyanobacteria, possess four EvoMining predictions and four antiSMASH hits. *scyB* is located next to this specialized metabolism branch. **c)** The majority of the *Pseudomonas* copies are labeled as conserved metabolism, with only two EvoMining predictions located in a divergent branch. *Pseudmonas* has a low average copy number per genome, which is reflected in almost every copy labeled as central metabolism. **d)** Archaea, the most expanded taxon, has a populated branch with expansions labeled as antiSMASH hits (cyan), but without any EvoMining prediction. The four lineages have MIBiG recruitments, but *scyB* is only shared between Actinobacteria, Cyanobacteria and Archaea.

In the following subsection we dissect the results provided by EvoMining for GDH by comparing phylogenetic trees obtained for each genomic lineage. In Archaea, GDH has on average 1.23 copies per genome while in Actinobacteria, Cyanobacteria and *Pseudomonas* this average is 0.74, 0.56 and 0.65, respectively. In the latter, GDH is part of the shell genome, **Table S1**. Thus, based in these observations, we specifically investigated the relationship between BGC, expansion rates and genomic lineages, and thus we expand our analyses to the ALS EF, which is one of the 42 conserved enzymes, also recruited by the scytonemin BGC (43) but not by the pactamycin BGC (42).

### EF from the same BGC may experience different evolutionary dynamics: the case of GDH and ALS

The enzyme GDH, distributed in all domains of life, catalyzes the reversible oxidative deamination of glutamate into alpha-ketoglutarate and ammonia. This EF exist in three classes according to their use of cofactors. The first class uses NAD+ and it is referred to as GDH(NAD+). The second class utilizes NADP+ and it is known as GDH(NADP+). And the third class uses both NAD+ and NADP+ and is therefore referred to as GDH(NAD+ & NADP+) (13). Moreover, GDH enzymes are very diverse and can be divided according to their taxonomic distribution and structural features (28). However, although this GDH classification reflects more closely in the evolutionary history of this enzyme, for the sake of simplicity we adopted for our analysis the three GDH classes according to their cofactor specificity. GDH(NAD+) is utilized for glutamate oxidation and GDH(NADP+) for fixing ammonia, although some enzymes from Archaea can perform equally well with both cofactors (13). NAD or NADP specificity has probably emerged repeatedly, as it has been shown that a few mutations can reverse specificity (44). This suggests that global sequence similarity does not indicate similar specificity, which is an important consideration when analyzing highly divergent EF from different genomic lineages.

A detailed examination of GDH EF showed that expansion events are not abundant neither in Actinobacteria, **Fig. 4a**, nor in Cyanobacteria, **Fig 4b**, and that they are practically absent from *Pseudomonas*, **Fig 4c**. In contrast, a significant number of expansions were found in Archaea, **Fig 4d**. Thus, we focused in Archaea, and performed a detailed annotation in order to make sense of the resulting GDH EvoMining tree, which was rooted with a seed sequence from *Sulfolobus*, predicted to be a dual NAD(P)+ utilizing enzyme (45). Notably, the three GDH classes alternate throughout the tree according to RAST annotation, **Fig S12**. As expected, most of the sequences classified as central metabolic enzymes were situated in the early or basal branches of the tree. A more divergent and larger clade, consisting almost exclusively of NAD(P) specific enzymes (46), included many expansions that are antiSMASH hits, with only two central metabolic enzymes. Functional annotation of the genomic vicinity of these GDH orthologues points towards a potential recruitment by specialized metabolism.

These recruitments were identified mainly in organisms from the genera *Haladaptatus, Haloterrigena, Natrialba, Natrinema, Natrialbaceae*, and *Natronococcus*, within a genomic context suggestive of the synthesis of terpenes, as it include enzymes related with geranyl pyrophosphate, a precursor to all terpenes and terpenoids (47). Lastly, despite its higher divergence, the following two major branches in the tree corresponded as well as enzymes devoted to central metabolism, **Fig 4d**.

In contrast with the broadly occurring GDH expansions related to metabolic adaptations in Archaea, the resulting Cyanobacteria’s tree showed expansions only in 4.5% of the genomes, **Fig 4b & Table S1**. Among these expansions, four antiSMASH and four EvoMining predictions conformed the branch that contain ScyB enzymes, which are GDH homologues. ScyB participates in the synthesis of scytonemin, a yellow sunscreen pigment produced by many Cyanobacteria to protect them against UV-A radiation (11,48,49). *Nostoc punctiforme* PCC 73102 is the scytonemin producer deposited at MIBiG. Unexpectedly, EvoMining only revealed a few GDH sequences from *Nostoc* species, even though *scyB* homologues can be found in their genomes, as it will be further discussed in the final phylogenomics subsection. This observation could be due to large sequence divergence between copies devoted to central and/or specialized metabolism in these organisms, despite being closely related.

Inspection of the genomic vicinity of the scytonemin BGC, furthermore, made us realized that the *scyB* gene is always next to the *scyA* gene, **Fig 5a**. This gene codes for a homologue of the acetolactate synthase large subunit, or ALS, an enzyme that was included amongst the 42 EF analyzed, with an average copy number of 1.87% in the entire Cyanobacteria database, and an average of 2.1 copies in organisms with at least a copy, but with a statistical mode of 1, **Table S1**.

**Figure 5.**
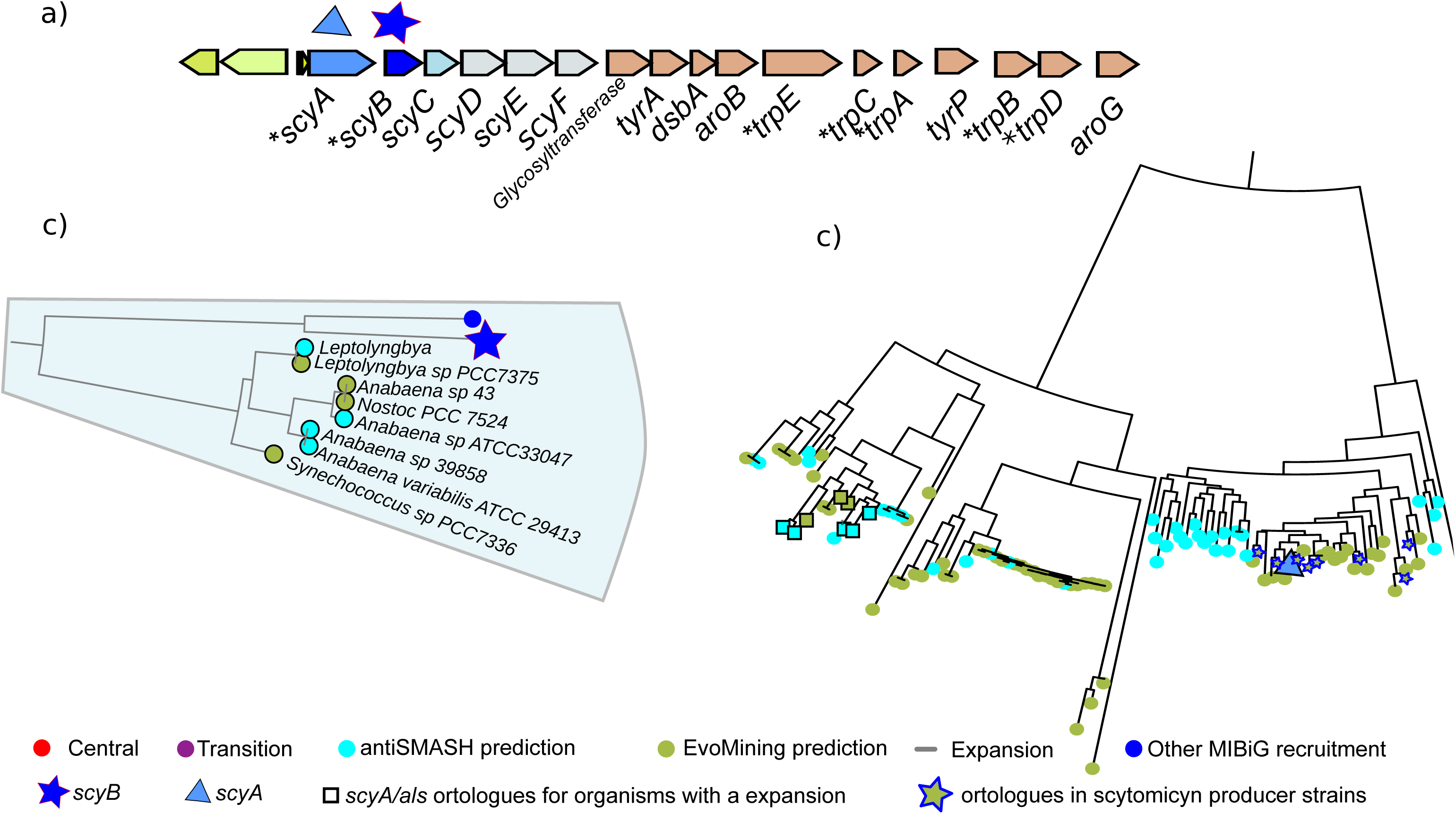
GDH and ALS recruitments into scytonemin BGC. **a)** Scytonemin BGC from *Nostoc punctiforme* is composed by regulatory genes (green), genes that participates in scytonemin biosynthesis (blue) and genes devoted to precursor supply (brown). Eight EF of the scytonemin BGC were found to have their origin within the 42 conserved EF, as shown by an asterisk. **b)** Zoom-in of Cyanobacteria GDH expansion branch close to *scyB*. Many of the known scytonemin producers are not found in this branch. **c)** Zoom-in of the *scyA* branch, showing ALS expansions correctly and exclusively marked by EvoMining with a fate in specialized metabolism. Known scytonemin producers are marked with stars. Squares indicate expansions devoted to specialized metabolism located in the genomic vicinity of GDH expansions that coincide with the *scyB* branch.

This data implies that many organisms have more than two ALS copies, which may correlate with the fact that this family showed larger dispersion around the mode, **Fig. S4 (blue line)**. After analysis of the Cyanobacteria ALS EvoMining tree, **Fig. S11**, it was found that *scyA* is in fact a recruitment localized in the same branch that contains ALS sequences from *Nostoc* spp, which were labeled as EvoMining predictions, **Fig 5c**. The latter predictions actually include more than twenty organisms that are known to be scytonemin producers (10,12,50), an observation that agrees with the fact that the sister branches in the ALS tree shows antiSMASH hits. Interestingly, organisms in this branch correspond to the same organisms revealed after the EvoMining analysis of GDH, which can be seen in the zoom-in of the ScyB branch, **Fig. 5b.**

These results together suggest co-evolution, via expansion-and-recruitment events, of ScyA and ScyB from ALS and GDH, respectively. However, it should be noted that the expansion rates of these EF were quite different. Indeed, the sequence similarity between the GDH EF and homologues of ScyB turned out not to be enough to reconstruct a ScyB branch with enough expansions to suggest the occurrence of an NP BGC. This contrasted with the scenario found when ALS and ScyA were analyzed. These results provide important lessons when using EvoMining as a genome-mining tool, as enzymes that co-evolve may be subject to different constraints and evolutionary rates.

### Phylogenomics analysis of the scytonemin BGC

The most characterized scytonemin BGC and its cognate pathway, shown in **Fig 5a** and **6**, consist of 18 genes (12,43). In addition to regulatory genes, this BGC includes the main biosynthetic genes, *scyABC;* tailoring-enzyme genes involved in late dimerization and oxidation steps, *scyDEF;* and in some cases, genes involved in precursor supply, namely, *tyrA, dsbA, aroB, trpE/G, trpC, trpA, tyrP, trpB, trpD, aroG*.

The enzymes TrpABCDEG and AroB are part of the aromatic amino acid and shikimic acid pathways, and they seem to provide precursors for the synthesis of scytonemin in the form of L-tryptophan and prephenate. Interestingly, ScyA and ScyB have an origin in central metabolism, as they are homologues of ALS and GDH, but they have evolved different substrate specificities, **Fig 6**. ALS joins two pyruvates, leading to S-2- acetolactate (14), while ScyB joins indole-3-pyruvate with p-hydroxy-phenyl-pyruvic acid. Analogously, but in central metabolism, GDH converts L-glutamate into 2-oxoglutarate (13), while ScyA catalyzes an oxidative deamination of tryptophan. The product of these two enzymes, acting sequentially, is a dipeptide, which is cyclized by ScyC. The pathway finishes with a series of oxidations and a dimerization step to yield scytonemin (11,48,51).

**Figure 6.**
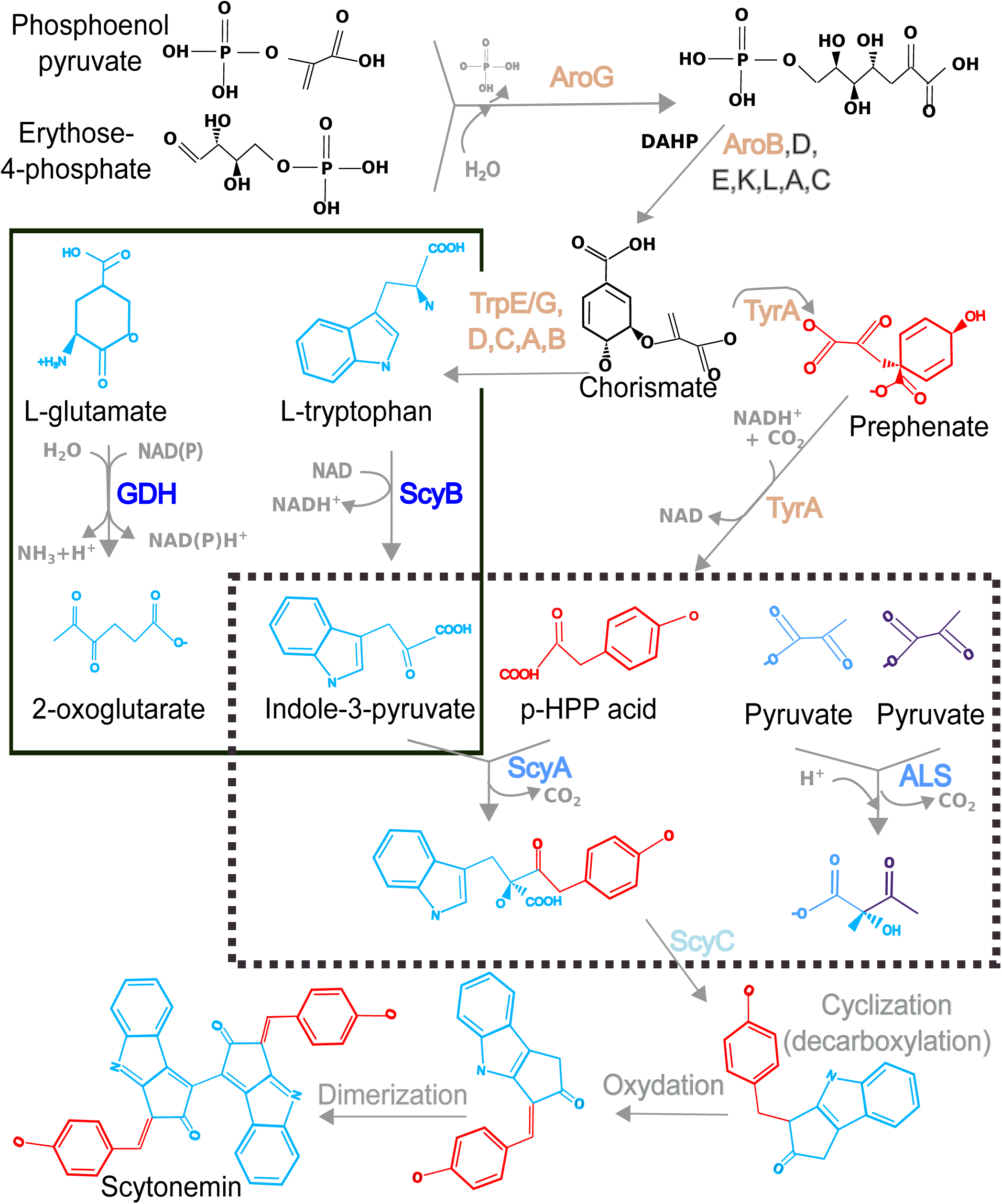
Metabolic origin and fate of GDH/ScyA and ALS/ScyB and scytonemin biosynthesis. AroG and AroB participates in chorismate synthesis, an internediary that is transformed in the precursors leading to ScyA substrates, i.e. L-tryptophan and prephenate. The reaction catalyzed by ScyB converting tryptophan into indole-3-pyruvate is similar to the conversion of L-glutamate into 2-oxoglutarate, catalyzed by GDH (solid square). ScyA catalyzes the decarboxylation of indole-3-pyruvate and p-hydroxy-phenyl-pyruvic acid (pHPP) acid to form a dipeptide that serves as scytonemin precursor, and, this reaction is analogous to the decarboxylation of two pyruvates by the original ALS enzyme (dotted square). ScyC performs a cyclization, and oxidation and dimerization steps conclude with scytonemin pathway. Enzymes in scytonemin BGC devoted to synthesis of precursors are colored in brown; scytonemin biosynthetic enzymes are colored blue.

In addition to GDH and ALS, seven of the EF that are present in the scytonemin BGC are part of the 42 EF analyzed herein. All of them have an origin in central metabolism and have been recruited into scytonemin BGC, **Fig 5 & Fig S5-S11**. From these, six of the seven EvoMining trees contain expansions that turned out to be EvoMining predictions, as the expansion branches include scytonemin genes, an indication of specialized metabolism. These families include AroB, as well as all genes in the L-tryptophan biosynthetic pathway, other than *trpF*.

The EvoMining predictions include enzymes from other sunscreen biosynthetic systems, such as shinorine and mycosporine-like aminoacids (52), **Fig. S10**, as well as other unrelated NP, including welwitindolinone (53), ambiguine (54) and fischerindoline (55), **Fig S5-S9**. These results illustrate how EvoMining can complement antiSMASH by identifying sequences that belong to nontraditional NP BGC, even when substrate specificity has not been changed.

To further investigate the presumed co-evolution of ScyA and ScyB we reconstructed the evolutionary history of these enzymes by concatenating their sequences, and contrasting the resulting phylogenetic reconstruction with the genomic vicinity of their cognate BGC, using CORASON (21). The resulting phylogenomic analysis revealed 34 cyanobacterial organisms with chemical diversity around their scytonemin BGC.

According to **Fig 7**, we could predict five additional putative chemical structures related to scytonemin, which correlates with gene losses-and-gains at these loci, **Fig 7**. The incorporation of genes that encode other enzymes, such as hydrolases (blue pattern), prenyltransferases (yellow pattern), and phosphodiesterases (purple pattern), could be forming compound **1**, a congener of scytonemin with a prenyl group. The predicted metabolite could be modified by the action of monooxygenases (pink pattern), which may reduce the carbonyl group to a hydroxyl group, to form compound **2**. Moreover, gene losses related to the enzymes ScyDEF might be driving the synthesis of compound **3**, an intermediary formed before the oxidation and dimerization steps. This intermediary could be modified by incorporation of a tyrosine moiety via a tyrosinase (gray pattern) and/or an amidase (orange pattern) to form compound **4**. We also found that *scyA* and *scyB* are part of a BGC that contains a hybrid NRPS-PKS (green and black pattern, respectively). Following biosynthetic logics related to these enzymes, we predicted compound **5**.

**Fig 7.**
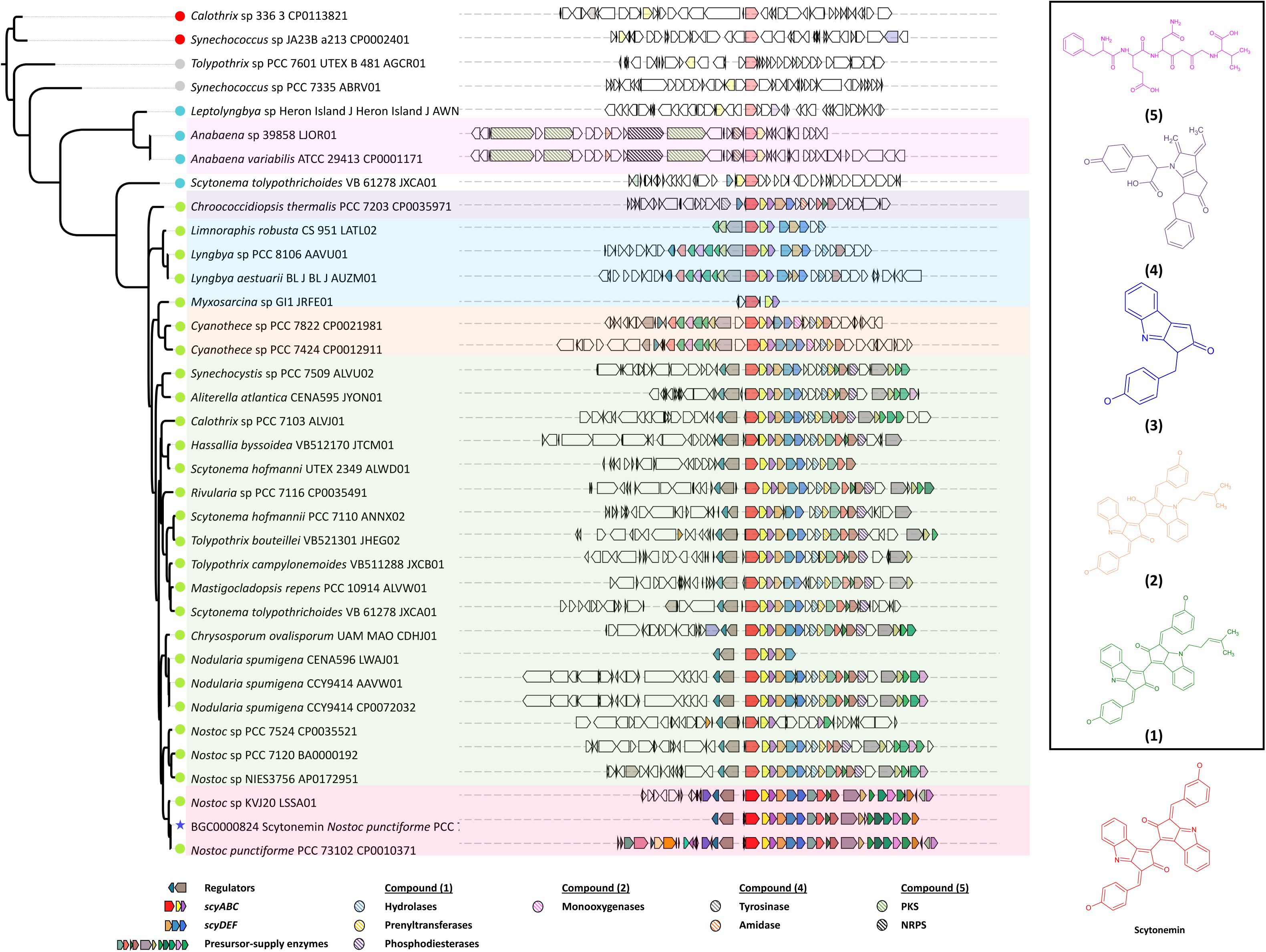
Phylogenomic analysis of *scyA* and *sycB* showed chemical diversity around the scytonemin BGC. Genomic vicinities containing both *scyA* and *scyB* in Cyanobacteria are shown next to a phylogenetic reconstruction using the protein sequences of these two genes. EvoMining classification for ALS family is shown in a circle. Relevant accessory genes in variants of scytonemin BGC, highlighted by colour boxes that define phylogenetic clades with biosynthetic signals, are label as follows: hydrolases (blue pattern), prenyltransferases (yellow pattern), phosphodiesterases (purple pattern), monooxygenases (pink pattern), tyrosinases (gray pattern) and amidases (orange pattern). The NRPS and PKS from the basal branches are colored in green and black patterns, respectively. Five putative chemical structures related to scytonemin were predicted considering biosynthetic logics and gain-and-losses of key genes, as shown in the hand-right panel of the figure.

The chemical diversity revealed by these predictions emphasizes the evolutionary dynamics of specialized metabolism, but which was only traceable by means of using ScyA and ScyB as beacons. These results demonstrate the increased predictive power of EvoMining for opening new metabolic spaces typically overlooked by standard NP genome mining approaches, which is potentialized when it is linked to other tools, such as CORASON.

### Final remarks and considerations for the use of EvoMining

EvoMining was developed as a stand-alone genome-mining tool and applied to selected Enzyme DB composed by EF common to highly divergent phyla. Our analyses lead to the conclusion that expansion-and-recruitment events are both EF and genomic-lineage dependent, an important consideration when using EvoMining. Although genome size seems to matter, we also found exceptions where EvoMining could predict novel BGC in relatively small genomes, suggesting that further analyses are needed to assess this relationship. Along these lines, we opted to compare genomic lineages that are not only highly divergent, and in some cases poorly understood with regards to NP biosynthesis, but also disproportionate in terms of their taxonomic resolution and distances. Thus, it is possible that these factors could have imposed a bias when establishing relationships between genome size, gene expansions and metabolic diversity.

After comprehensively analyzing GDH, an EF notably expanded in Archaea but not in other taxa, we provided an example of a recruitment of a central metabolic enzyme into a NP BGC, as well as into other metabolic pathways. It is interesting to note that the most expanded EF in previous EvoMining proof-of-concept analyses (15) were asparagine synthase, 2-dehydro-3-deoxyphosphoheptanoate aldolase and 3-phosphoshikimate-1- carboxivinyl transferase, which lead to the discovery of unprecedented arsenolipid biosynthetic enzymes. Notably, none of these enzymes were part of the 42 EF analyzed herein, reinforcing the notion that not only conserved enzymes, but also shell enzymes with extra copies, can serve as beacons for the discovery of novel NP BGC. These observations emphasize the predictive nature of EvoMining, but which became only apparent after the origin and fate of enzymes could be traced back to evolutionary events at different levels, from genome dynamics involving large loci, to different mutations rates at the protein sequence level.

EvoMining users, therefore, should define beforehand the most appropriate EF to be used for a certain taxonomic group. Selected Enzyme DB should contain a set of EF where expansion patterns could be detected. In turn, EF with a distribution restricted to a small percentage of genomes are not suitable for EvoMining analysis. It is also important to determine which EF are shared by most of the genomes in the genomic lineages of interest, and whether or not this is important for the type of EvoMining analyses to be performed. Original EvoMining DB included manually curated EF only involving central metabolic enzymes, but as it was demonstrated here these did not necessarily represent the core enzymatic repertoire of Actinobacteria. This relates to the difficulty of defining what is central metabolism, and thus we prefer to use the term core enzymes at different thresholds of conservation, being 50% the one used to define shell enzymes. This notion implies the possibility of automatizing Enzyme DB integration by selecting for EF in any given genomic lineage, avoiding the need to arbitrarily define what is central metabolism.

Another key point in EvoMining success relates to the improvement of the NP DB due to the availability of MIBiG (2). The previous EvoMining version did not include a cyanobacterial NP BGC, and for this reason only this EvoMining version could identify hits for the ScyA and ScyB branches. Nevertheless, in absence of the signals provided by MIBiG, the extra copies of these EF would have been marked by EvoMining as expansions not involved in central metabolism. This maybe the case in Archaea, where some sequences in the GDH tree are labeled as such, possibly related to terpenes as well as to other metabolic fates yet-to-be discovered. The poor representation of Archaea at MIBiG is clearly due to the little investigation available in the potential of Archaea to synthesize NP, as our results suggest that current methods based in previous knowledge from unrelated taxa impose biases that hamper our ability to unlock the metabolic diversity of this domain of life. We anticipate that this situation will be overcome by EvoMining, as it is a less-biased and rule-independent approach.

## Acknowledgements

We would like to thank CONABIO, Mexico, for access to computing facilities. We express our gratitude to Jose Daniel Hernandez, Ernesto Campos and Alan Martínez Guerrero for their technical support. We also thank Lianet Noda, Karina Verdel and Adriana Espinosa for useful discussions that helped shaping this manuscript, and Pablo Cruz for intellectual insights and for a sustained and open scientific dialogue.

## ABBREVIATIONS

BGC: Biosynthetic Gene Clusters
NP: Natural products
GDH: Glutamate dehydrogenase
ASL: Acetolactate synthase large subunit
EF: Enzyme families
DB: database
BBH: Bidirectional Best Hits

